# A combinatory genetic strategy for targeting neurogliaform neurons in the mouse basolateral amygdala

**DOI:** 10.1101/2024.01.03.573867

**Authors:** Attila Ozsvár, Meike Claudia Sieburg, Monica Dahlstrup Sietam, Wen-Hsien Hou, Marco Capogna

**Author notes:** **Correspondence:** Wen-Hsien Hou. Deceased (Marco Capogna).

## Abstract

The mouse basolateral amygdala (BLA) contains various GABAergic interneuron subpopulations, which have distinctive roles in the neuronal microcircuit controlling numerous behavioral functions.

In mice, roughly 15% of the BLA GABAergic interneurons express neuropeptide Y (NPY), a reasonably characteristic marker for neurogliaform cells (NGFCs) in cortical-like brain structures. However, genetically labeled putative NPY-expressing interneurons in the BLA yield a mixture of interneuron subtypes besides NGFCs. Thus, selective molecular markers are lacking for genetically accessing NGFCs in the BLA.

Here, we validated the NGFC-specific labeling with a molecular marker, neuron-derived neurotrophic factor (NDNF), in the mouse BLA, as such specificity has been demonstrated in the neocortex and hippocampus. We characterized genetically defined NDNF-expressing (NDNF+) GABAergic interneurons in the mouse BLA by combining the Ndnf-IRES2-dgCre-D transgenic mouse line with viral labeling, immunohistochemical staining, and in vitro electrophysiology.

We found that BLA NDNF+ GABAergic cells mainly expressed NGFC neurochemical markers NPY and reelin (Reln) and exhibited small round soma and dense axonal arborization. Whole-cell patch clamp recordings indicated that most NDNF+ interneurons showed late spiking and moderate firing adaptation. Moreover, ∼81% of BLA NDNF+ cells generated retroaxonal action potential after current injections or optogenetic stimulations, frequently developing into persistent barrage firing. Optogenetic activation of the BLA NDNF+ cell population yielded both GABA_A_- and GABA_B_ receptor-mediated currents onto BLA pyramidal neurons (PNs). We demonstrate a combinatory strategy combining the NDNF-cre mouse line with viral transfection to specifically target adult mouse BLA NGFCs and further explore their functional and behavioral roles.

## Introduction

The amygdala is an evolutionarily conserved brain structure located deeply in the temporal lobe (Janak and Tye, 2015). In rodents, the amygdala can anatomically be divided into the basolateral and the central sectors (Pitkänen et al., 1997; Ledoux, 2004). The basolateral amygdala complex (BLA) is critical for processing information related to reward, valence, and emotional behaviors such as fear and anxiety (Ledoux, 2012; Calhoon and Tye, 2015). The central sector comprises the central amygdala and intercalated cell masses, which are thalamic-like structures comprising local- and long-range projecting GABAergic interneurons (Capogna, 2014). The central amygdala is key to diverse behaviors such as anxiety, reinforcement, social interaction, and survival actions such as fear expression and cataplexy (Janak and Tye, 2015; Pape and Pare, 2010; Moscarello and Penzo, 2022; Mahoney et al., 2017).

The BLA receives major afferents from the sensory thalamus and cortices, ventral hippocampus, medial prefrontal cortex, nucleus accumbens, and extended amygdala regions (Ledoux, 2000; Pare and Duvarci, 2014). It is a cortical-like structure comprising 80% excitatory glutamatergic PNs and 20% inhibitory GABAergic interneurons (Gaudreau and Pare, 1996; Pape and Pare, 2010; Vereczki et al., 2021; Hajos, 2021). The latter population comprises heterogeneous clusters characterized by morpho-electrophysiological and neurochemical properties and subcellular targeting areas of the postsynaptic cells (Vereczki et al., 2020; Hájos, 2021). The activity of PNs is shaped by phasic and tonic inhibitions generated by diverse GABAergic interneuron populations (Farrant and Nusser, 2005; Ehrlich et al., 2009; Wolff et al., 2014). Phasic inhibition onto PNs can either originate from perisomatic areas (Veres et al., 2014; Veres et al., 2017) or dendritic areas (Krabbe et al., 2018; d’Aquin et al., 2022), while tonic inhibition tunes the offset control of PN spike probability through extrasynaptic GABAA- and GABAB receptors (GABAAR and GABABR, respectively; Marosky et al., 2004, Manko et al., 2012).

One major source of tonically released GABA in BLA is the activation of neurogliaform cells (NGFCs, Manko et al., 2012; Rovira-Esteban et al., 2019). In the mouse BLA, NGFCs represent about 15% of the total GABAergic interneuron populations (Vereczki et al., 2021). Similar to observations made in the hippocampus and neocortex, BLA NGFCs have small, spherical soma and short, repeatedly branching dendrites primarily confined in the dense axonal arborization (Manko et al., 2012; Overstreet-Wadiche and McBain, 2015). Despite the relatively small occupied volume of the axonal arborization, its presynaptic bouton density is noticeably high along the axon processes (Olah et al., 2009; Ozsvar et al., 2021). Such a unique structural feature promotes unparalleled signaling capability to reach not only postsynaptic GABAAR but reliably activate extrasynaptic GABAAR as well as GABABR, triggering a biphasic mixture of fast and slow inhibition on target neurons (Tamas et al., 2003; Manko et al., 2012).

Consistent with results from other cortical-like structures, NGFCs in the mouse BLA mainly express neuropeptide Y (NPY) and low levels of cholecystokinin (CCK) and somatostatin (Sst, Manko et al., 2012; Rovira-Esteban et al., 2019; Vereczki et al., 2021). Such complex molecular profiling makes it challenging to isolate NGFCs from other BLA interneuron types by approaches based on classical interneuron molecular markers (Rovira-Esteban et al., 2019; Vereczki et al., 2021; Hajos, 2021). Although the BLA interneuron types have been more extensively characterized in the last decade (Wolff et al., 2014; Capogna, 2014; Verseki et al., 2021), no available combination of techniques has made it possible to target NGFCs in the mouse BLA selectively. Hence, how BLA NGFCs contribute to the circuit operation and their vulnerability to behavioral disorders remain unknown. Recent studies have described a genetic marker named neuron-derived neurotrophic factor (NDNF) to effectively target NGFCs in the neocortex and hippocampus (Tasic et al., 2016; Abs et al., 2018; Mercier et al., 2022). This study proposes and validates a strategy to selectively identify BLA NGFCs in mice by combining the Ndnf-IRES2-dgCre-D mouse line with GABAergic enhancer mdlx-dependent viral transfection.

## Materials and Methods

### Experimental animal model and subject details

The Ndnf-IRES2-dgCre-D mice (hereafter NDNF-cre, Tasic, et al., 2016) were purchased from the Jackson laboratory (stock #028536) and bred in the animal facility of Aarhus University. All mice were bred onto the C57BL/6J genetic background. Mice of both sexes (postnatal 8-13 weeks) were used for the experiments. Mice were given food and water ad libitum and group-housed on a 12 h light/dark cycle. All animal experiments were performed according to standard ethical guidelines and were approved by the Danish National Animal Experiment Committee (Permission No. 2017−15−0201−01201).

### Viral vectors

For labeling NDNF+ GABAergic neurons in the mouse BLA, we injected ssAAV-DJ/2-mDlx-HBB-chI-dlox-EGFP(rev)-dlox-WPRE-bGHp(A) (6×10^12 vector genomes/ml, v241-Dj, VVF Zurich, Switzerland). In a subset of experiments, ssAAV-DJ/2-mDlx-HBB-chI-dlox-hChR2(H134R)-mCherry(rev)-dlox-WPRE-bGHp(A) (6×10^12 vector genomes/ml, v317-Dj, VVF Zurich, Switzerland) was used for optogenetic assisted target cell mapping.

### Stereotaxic injection

Mice (NDNF-cre, n = 25, 6-10 weeks old) were anesthetized using FMM: a mix of 0.05 mg/ml Fentanyl (0.05 mg/kg, Hameln), 5 mg/ml Midazolam (5 mg/kg, Hameln) and 1 mg/ml Medetomidine (0.5 mg/kg, VM Pharma). Mice were placed onto the stereotaxic frame (Kopf instruments) and maintained at a constant body temperature (34-36°C). A midline scalp incision (∼0.8 cm) was made, and small craniotomies were made to target BLA bilaterally (coordinates from bregma: AP: −1.47 mm, ML: ±3.4 mm, DV: −4.45, -4.7 mm). Coordinates were normalized to a bregma-lambda distance of 4.21 mm. A total of 250 nl/site of viruses were injected through a glass capillary (Harvard Apparatus) by pulses of the Picospritzer III (Parker Hannifin). The pipette was raised 0.1 mm for an additional 10 min to minimize the upward flow of viral solution and was slowly withdrawn. After the viral injection, the incision was closed by suturing. Mice were given an antidote mix of 0.4 mg/ml Naloxone (1.2 mg/kg, B. Braun), 5 mg/ml Atipamelozone Hydrochloride (Revertor; 2.5 mg/kg, Vibrac AG), and 0.5 mg/ml Flumazenil (0.5 mg/kg, Sintetica).

### Acute brain slice preparation and patch-clamp recording

Mice were anesthetized with isoflurane (Zoetis) and decapitated. The brains were removed, and 300 µm-thick coronal sections were prepared by a vibratome (Leica) using ice-cold sucrose-based artificial cerebrospinal fluid solution (ACSF, Ozsvar et al., 2021) containing (in mM) 84 NaCl, 25 NaHCO3, 1 NaH2PO4, 2.5 KCl, 25 glucose, 75 sucrose, 0.5 CaCl2, and 4 MgCl2. Slices were recovered in an oxygenated (95% O2 and 5% CO2) chamber containing sucrose and saline at 34°C for 30 min and then kept at 18°C. During the experiment, slices were transferred to a submerged chamber and perfused with oxygenated ACSF containing (in mM) 130 NaCl, 24 NaHCO3, 1 NaH2PO4, 4 KCl, 10 glucose, 1.3 CaCl2, and 0.9 MgCl2. A pE-300 (CoolLED) was used in a subset of experiments to provide optogenetic stimulations (1 mW). 10 μM gabazine (Tocris Bioscience) and 10 μM CGP-55845 (Tocris Bioscience) were added to the ACSF to isolate GABA_A_R and GABA_B_R-dependent responses.

The recorded cells’ epifluorescence expression and soma location were visually confirmed and selected under an infrared differential interference contrast (IR-DIC) CCD camera (Scientifica). Whole-cell patch-clamp recordings were made with a Multiclamp 700B amplifier (Molecular Devices). Recording electrodes (3–6 MΩ) were pulled from borosilicate glasses (outer diameter, 1.5 mm; 0.32 mm wall thickness; Harvard Apparatus) and filled with internal solution containing (in mM): 126 K-gluconate, 4 KCl, 4 Mg-ATP, 10 HEPES, 10 phosphocreatine, 0.3 Na-GTP, with or without 0.2 % biocytin (Thermo Fisher Scientific) with pH adjusted to 7.3 with KOH. The series resistance (Rs) was compensated to 70-80% in the voltage-clamp configuration. The data were discarded if Rs > 20 MΩ or Rs change > 20 % throughout the recording. Signals were low-pass filtered at 4 kHz (four-pole Bessel) and sampled at 10 kHz using a digitizer (Digidata 1440A; Molecular Devices).

### Immunohistochemistry (IHC)

For characterizing the molecular identities of the eGFP+ cells, virally injected NDNF-cre mice were anesthetized by FMM injection (I.P.) and perfused with ice-cold PBS plus heparin (50 mg/ml) followed by 4% PFA. Fixed brains were removed, post-fixed in 4% PFA for 1 hr, and stored in 0.1M PBS. Brains were sliced into 60 or 100 μm coronal sections using a vibratome (Leica 1000S, Leica). Collected brain slices were washed with PBS and blocked in a buffer containing 10% normal donkey serum (NDS), 10% normal goat serum (NGS), 1% bovine serum albumin (BSA), and 0.5% PBST for 6 hr at 4°C. After blocking, slices were incubated with primary antibodies: Chicken Anti-GFP (1:1000, ab13970, Abcam), Rabbit Anti-NPY (1:3000, #22940, ImmunoStar), Rabbit Anti-Reln (1:1000, ab230820, Abcam) or Mouse Anti-Reln (1:1000, MAB5364, Merck) diluted in a solution containing: 1% NDS,1% GDS, 3% BSA, and 0.5% PBST for 48 hr at 4°C. After washing with 0.5% PBST, slices were incubated with secondary antibodies: Alexa 488 Donkey Anti-Chicken (1:400, AB_2340375, JacksonImmuno), Alexa 568 Donkey Anti-Rabbit (1:400, A10042, Invitrogen) diluted in antibody solution ON at 4°C. Slices were washed in 0.5% PBST and mounted onto gelatin-coated slides with DAPI-containing mounting medium (FluoromountG, Vectashield) and coverslipped. Images were taken as tile z-stacks under a 20x objective by an IX83 (Olympus) spinning-disk confocal microscope at 5 µm step size.

### Morphological recovery of biocytin-filled cells

During recordings, eGFP+ neurons (n = 21 cells) were filled with biocytin (0.2%). After ∼15 min of recording, slices were fixed overnight with 4% PFA in PBS (0.1 M, pH 7.3). After washing with Alexa594-conjugated streptavidin (2 µl/ml; Invitrogen) in PBS containing 0.3% PBST for 2 hr at 4°C. After washing, slices were mounted onto gelatin-coated slides with DAPI-containing mounting medium (FluoromountG, Vectashield) and coverslipped.

Recovered neurons (12/21 cells) were examined by an LSM 980 Airyscan 2 (Zeiss) confocal microscope using a 40x 1.4 NA oil immersion objective. The morphology of the cells was reconstructed from a stack of 120-202 images per cell. Image stacks belonging to a single cell were imported into the Neuromantic 1.7.5 software (Myatt et al., 2012) for 3D reconstruction.

### Data analysis and statistics

The electrophysiological data were recorded with pCLAMP 10.6 software (Molecular Devices). The electrophysiological recordings were analyzed by custom-written Matlab scripts (MathWorks). The properties of action potentials (APs) were analyzed from the first suprathreshold voltage responses triggered by depolarizing current injection steps (1 s). The AP peak, width, and afterhyperpolarization (AHP) amplitude were calculated from the onset point of the AP. The AP onset was determined by the projection of the intersection of two linear fits. The first line was fitted to the baseline 2 ms window, and the second line was fitted to the 5-30% of the rising phase of the AP. The rate of depolarization was measured as the linear fit of the trace 1 ms before the onset. The spike latency was calculated on rheobase traces with at least two APs by measuring the delay between the first AP onset time. The input resistance (Rin) was determined from the slope of a linear fit to the subthreshold voltage responses vs. current injection, where stimulating current ranged from -110pA to +30pA. The membrane time constant (**τ**) was calculated by fitting a single exponential to the voltage response from -110 pA current injection. The membrane capacitance was calculated as **τ**/Rin. The adaptation index was calculated as the ratio of the first and last inter-spike interval, measured at the first suprathreshold voltage response. For the analysis of the retroaxonal barrage firing data, APs were detected by their pea. Statistic tests were performed and plotted using Prism 6.0 (GraphPad Software). The data distributions were examined by the d’Agostino-Pearson normality test. Statistical significance was tested by the Mann-Whitney U (MWU) test or Wilcoxon signed-rank (WSR) test, and the significance level (p) indicated. Data were presented as mean ± the standard error of the mean (SEM) unless otherwise stated. Significance levels were set at p < 0.05. ** denotes p< 0.01, *** denotes p< 0.001.

## Results

### NDNF+ GABAergic neurons in the mouse BLA

As NDNF has been proposed as an NGFC-specific genetic marker in the mouse neocortex and hippocampus (Abs et al., 2018; Tasic et al., 2018; Mercier et al., 2022), we aimed to determine if the NDNF-cre mouse line could be used to identify the NGFCs in other cortical-like structures such as BLA. Since there is a likelihood of a low-level Cre expression in PNs during development of the NDNF-cre mouse line (Guo et al., 2021; Mercier et al., 2022), we injected a recombinant adeno-associated viral vector rAAVdj-mdlx-DIO-eGFP carrying an enhanced green fluorescence protein driven by mdlx enhancer (Dimidschstein et al., 2016) in the BLA of NDNF-cre mice (Fig.1A). Two weeks later, we examined the eGFP expression pattern and found that the combinatory strategy yielded a sparse labeling pattern (10 - 30 cells per 100-μm section, Fig.1A). We further tested if the eGFP+ cells within the BLA expressed classical NGFC markers such as NPY and Reelin (Reln) by double and triple IHC (Fig.1B). We observed that most BLA eGFP+ cells express either NPY (81.02 ± 2.15%; Fig.1C) or Reln (79.75 ± 2.96%; Fig.1D) in double IHC experiments. In a subset of triple IHC experiments, we found that most BLA eGFP+ cells coexpress Reln and NPY (68.83%, Fig.1E), and only a minor fraction of cells that are negative to both NPY and Reln (11.82%, Fig.1E). Moreover, we did not observe eGFP+ cells with PN-like morphology. These results indicate that the BLA cells labeled by this combinatorial strategy are likely GABAergic interneurons and correspond to one major uniform cell population with similar neurochemical identities.

**Figure 1.**
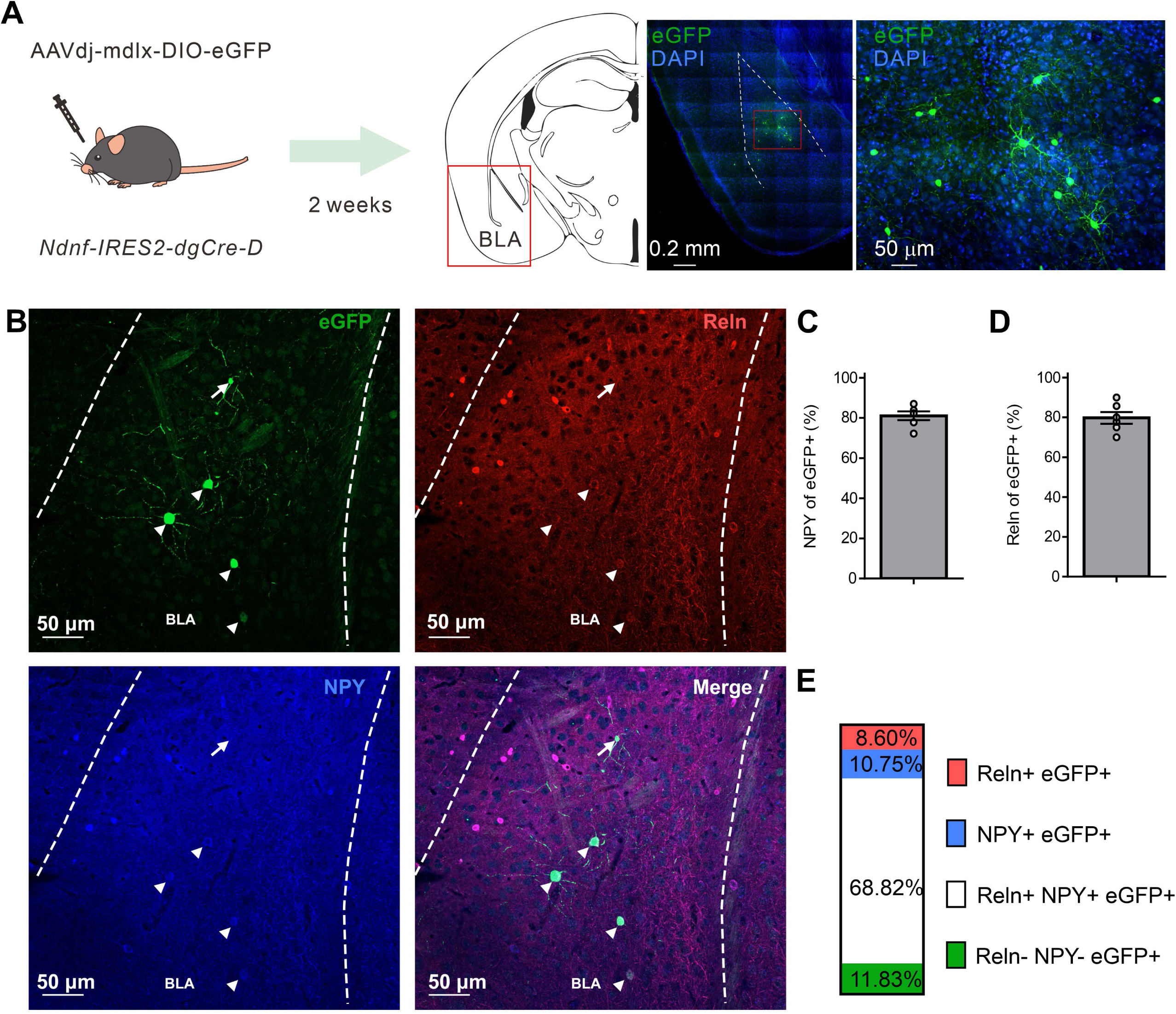
NDNF-expressing (NDNF+) GABAergic neurons in the mouse BLA. **(A)** Left, the scheme of genetic strategy; middle, a representative tile image of the amygdala region with eGFP amplification and DAPI staining; right, an enlarged view from the indicated red rectangular area from the middle image. (B) Confocal immunofluorescence micrographs showing amplified eGFP, NPY, and Reln expressions in the BLA. Arrowheads indicate triple immunopositive neurons, while the arrow depicts an eGFP-only cell. (C) Quantification bar graph of eGFP colocalization with NPY or (D) Reln (n = 6 sections, 3 mice, mean ± SEM). (E) Quantification bar graph of eGFP colocalization with NPY and Reln (n = 6 sections, 2 mice).

### Morpho-electrophysiological characterization of BLA NDNF+ GABAergic cells

Next, we wondered if the physiological and morphological properties of the virally labeled BLA NDNF+ GABAergic interneurons mainly corresponded to NGFCs. We performed whole-cell recordings from acute mouse brain slices and biocytin-assisted *post hoc* morphological identification. The recorded neurons had small round or elongated somata, fine, short dendrites, and extensively branching axonal processes containing numerous varicosities (Fig.2A-C).

**Figure 2.**
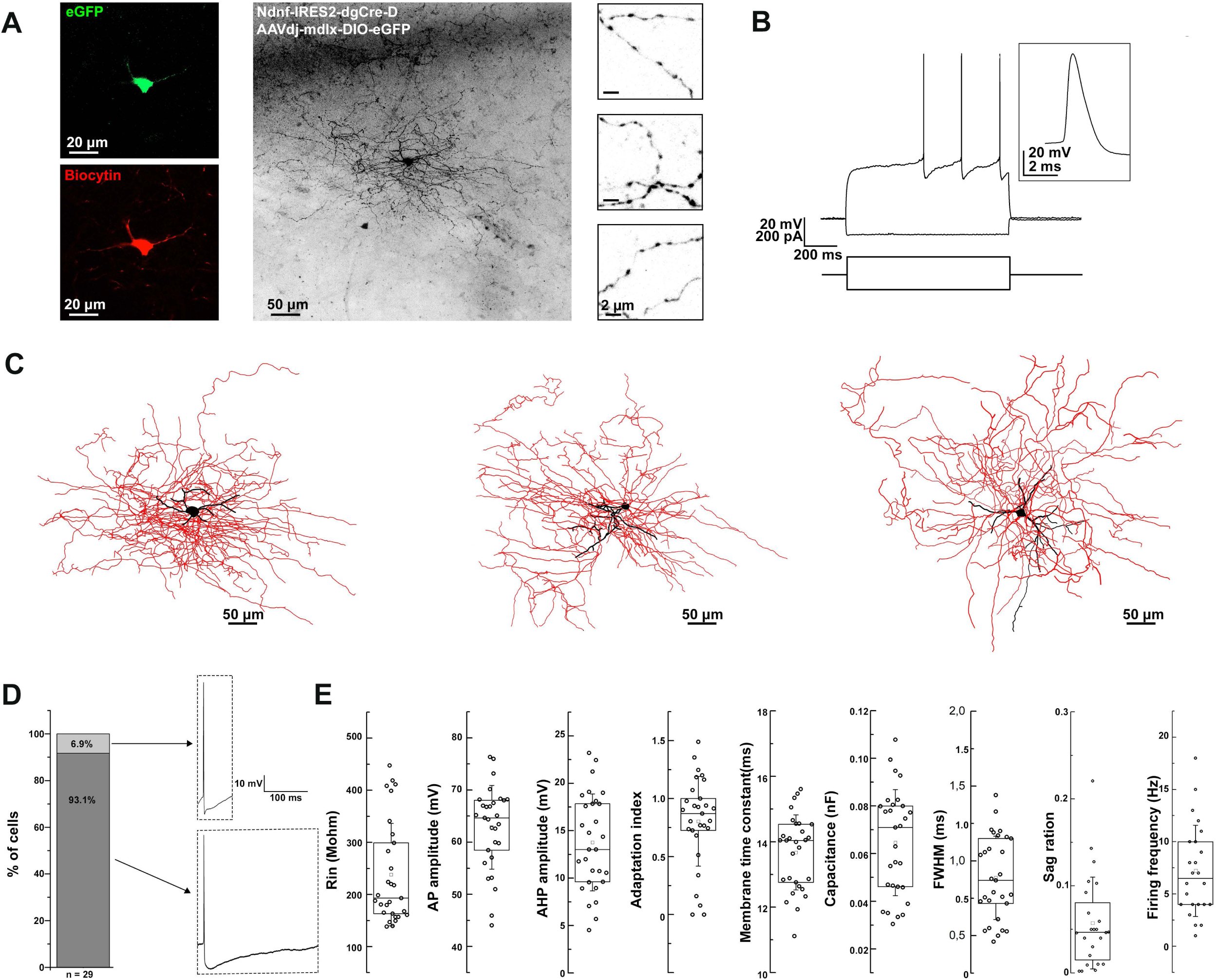
Morpho-electrophysiological property characterization of BLA NDNF+ GABAergic cells. **(A)** Confocal image stacks of a biocytin-labeled eGFP+ cell in the BLA of an NDNF-cre mouse. Confocal immunofluorescence micrographs showing immunohistochemically amplified eGFP and biocytin signals. On the right side, high-magnification insets show axonal segments with dense boutons. Scale bars, 20, 50, and 2 µm. **(B)** Passive and active electrophysiological responses from the same eGFP+ cell. Voltage responses to long-pulse current injections (-110 pA and +170 pA, 1 s). Inset, a single AP evoked by a short current pulse (+180 pA, 2 ms) recorded from the same cell. (C) Three-dimensional reconstruction of the same cell (left), and two additional reconstructions from separate experiments (middle and right). Black color represents the soma and dendrites, while red represents the axon. Scale bars 50 µm. (D) Left, a bar graph indicating that only a small fraction of BLA NDNF+ GABAergic interneurons exhibit non-NGFC AP properties such as sharp AHP peak and fast repolarization kinetics (n = 2 out of 29 cells). Right, representative APs from non-NGFC-like (top) and NGFC-like cells (bottom). (E) Box plots representing the different passive and active electrophysiological parameters from all recorded BLA NDNF+ GABAergic interneurons (n = 29 cells, n = 21 mice). For the boxplots, boxes represent the interquartile range, small squares at the center represent the mean, horizontal lines indicate the median, and whiskers show the SD.

In terms of electrical properties, the vast majority of the recorded eGFP+ cells exhibited characteristic features such as a slow depolarizing ramp and following late spiking phenomenon similar to previously described BLA NGFCs (Fig.2B; Manko et al., 2012). These cells had relatively low input resistance and fast membrane time constant. The action potentials (APs) were relatively broad, followed by a pronounced AHP (Fig.2D). The firing patterns were mostly moderately adapting during suprathreshold depolarization, with a relatively low firing frequency. Upon hyperpolarizing current injection, the recorded cells displayed a small sag ratio, indicating low/lack of h-current (Fig.2E and Table1). Only a minor fraction of the recorded cells exhibited sharp AHP peak and fast repolarization kinetics, compared to the vast majority displaying remarkably slow repolarization kinetics (Fig.2D). In conclusion, the electrophysiological features of the virally labeled BLA NDNF+ GABAergic interneurons resemble remarkably similar features to previously described BLA NGFCs.

### Retroaxonal barrage firing in BLA NDNF+ GABAergic interneurons

NGFCs in the neocortex and hippocampus exhibit persistent firing upon continuous depolarization (Suzuki et al., 2014; Chittajallu et al., 2020). We tested whether BLA NGFCs exhibited similar persistent firing properties in some experiments. We prepared acute brain slices and performed whole-cell patch-clamp recordings from virally labeled NDNF+ cells in BLA using NDNF-cre mice injected with AAVdj-mdlx-DIO-eGFP. The persistent firing was induced using repetitive positive current injection steps (+300 pA, 1 s, Fig.3B). We induced the barrage firing in 13 of 17 tested cells. Phase plots revealed that triggered APs and retro-axonal firing action potentials (rAPs) had two components: an initial component represented spiking in the axon and a second component that overlapped with the current-evoked spikes, indicative of a somato-dendritic spike following the initial, axonally initiated spike (Fig.3C). Further analysis revealed distinct characteristics in the subthreshold rate of depolarization and the AP threshold of rAPs compared to spikes evoked by somatic current injection (Rózsa et al., 2023). The rAPs during the persistent firing showed a lower subthreshold rate of depolarization compared to APs during repetitive current injection steps (0.99 ± 0.65 vs. 1.9 ± 0.67 mV/ms, mean ± SD, WSR-test, p < 0.001; n = 8 cells, Fig.3D-E). During persistent firing, the rAPs exhibited a more hyperpolarized AP threshold than somatic APs (population: -58.3 ± 2.34 vs. -42.13 ± 2.43 mV, WSR-test, p < 0.001, Fig.3F). The subthreshold rate of depolarization and threshold of the APs proved to reliably separate the somatic and axon-initiated APs (Fig.3G).

**Figure 3.**
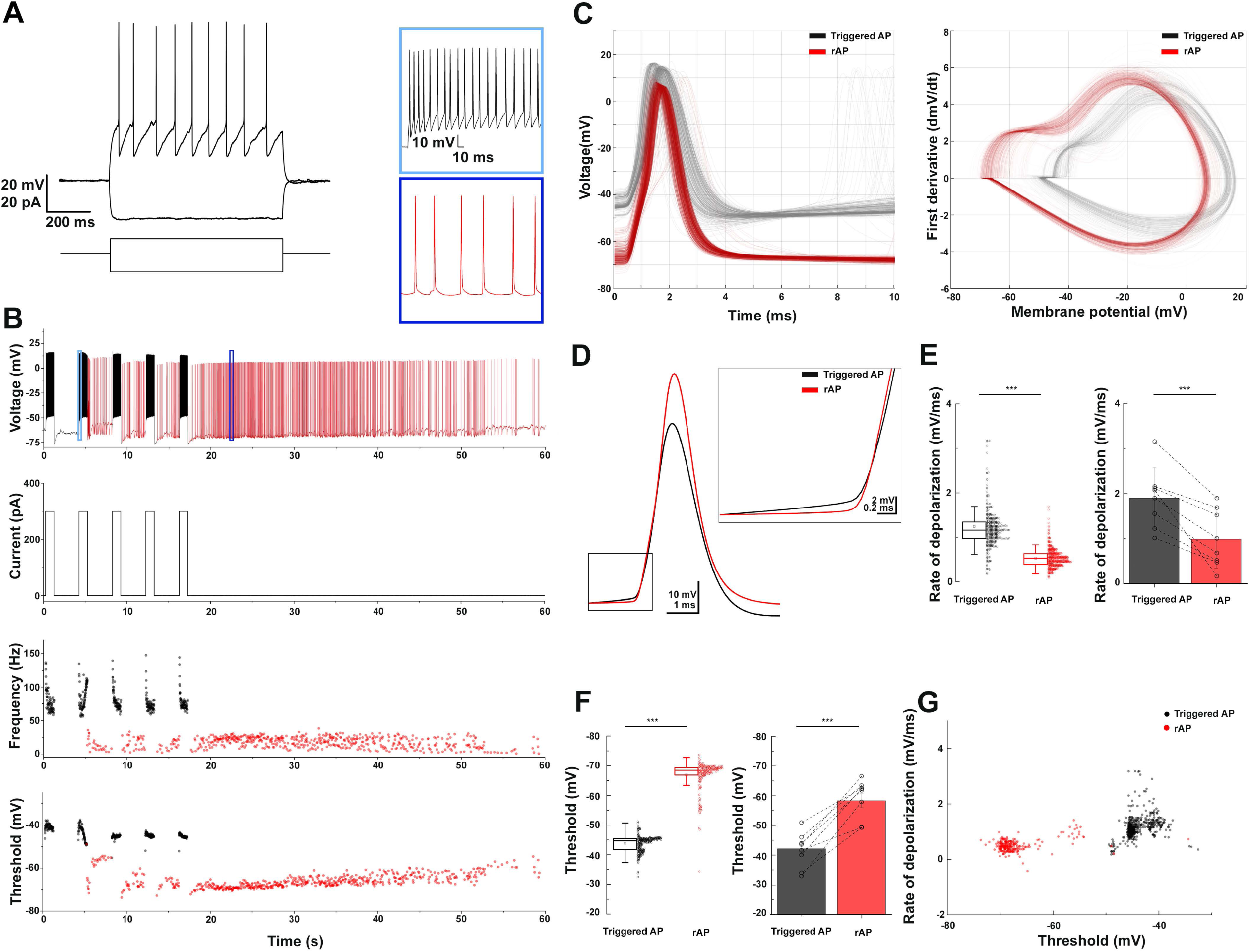
Retroaxonal barrage firing in BLA NDNF+ GABAergic neurons. **(A)** The representative firing pattern of a recorded NDNF+ interneuron in the BLA. **(B)** The induced retroaxonal barrage firing after repetitive suprathreshold current injection (+300 pA, 1 s). Light and dark blue boxes show enlarged sections of triggered and retroaxonal barrage firing patterns. The current injection protocol is indicated below. Scatter plots show the frequency and AP threshold throughout the stimulation and retroaxonal barrage firing period from a single cell. Current injection-triggered APs are presented in black, and spontaneous rAPs are depicted in red. (C) Triggered APs and rAPs recorded from the single cell shown on panel (B). Left, superimposed triggered APs (black) and rAPs (red). Right, phase plot of triggered APs (black) and rAPs (red). (D) Averaged traces of the triggered AP (black) and rAP (red) from a single cell show distinct rates of depolarization before the AP onset. (E) Left, a box plot of the triggered APs vs. rAPs depolarization rate values from a single cell (1.22 ± 0.71 vs. 0.5 ± 0.21 mV/ms, mean ± SD, MWU-test, p < 0.001); Right, a bar plot shows the population average (1.9 ± 0.67 vs. 0.99 ± 0.65 mV/ms, mean ± SD, WSR-test, p < 0.001; n = 8 cells). (F) Left, a box plot of the triggered APs vs. rAPs threshold values from a single cell (-43.84 ± 2.59 mV vs. -66.85 ± 4.87 mV, mean ± SD, MWU-test, p < 0.001); Right, a bar plot shows the population average (-42.13 ± 2.43 mV vs. -58.3 ± 2.34 mV, mean ± SD, WSR-test, p < 0.001). For the boxplots, boxes represent the interquartile range, small squares at the center represent the mean, horizontal lines indicate the median, and whiskers show the SD. (G) A 2D scatter plot showing clear segregation of triggered APs and rAPs using AP threshold and rate of depolarization from a single cell.

### Functional assessment of BLA NDNF+ GABAergic cells by optogenetic

NGFCs have been reported in different brain areas to provide distinctive biphasic GABA_A_R and GABA_B_R-mediated synaptic inhibition with slow kinetics on target cells (Price et al., 2005; Manko et al., 2012). To confirm the synaptic output of BLA NDNF+ cells, we expressed Channelrhodopsin-2 (ChR2) in BLA NDNF+ cells by injecting AAVdj-mdlx-DIO-ChR2-mCherry in the BLA of NDNF-cre mice. After 3-4 weeks, we first tested the ChR2 expression by performing whole-cell recordings on the epifluorescence-identified mCherry+ cells (Fig. 4B). The mCherry+ cells also demonstrated similar NGFC-like morphology as previously presented, and brief light pulse trains (5 ms, 10 Hz) reliably triggered APs in mCherry+ cells (3 of 3 tested cells, n = 2 mice; Fig.4B-E).

**Figure 4.**
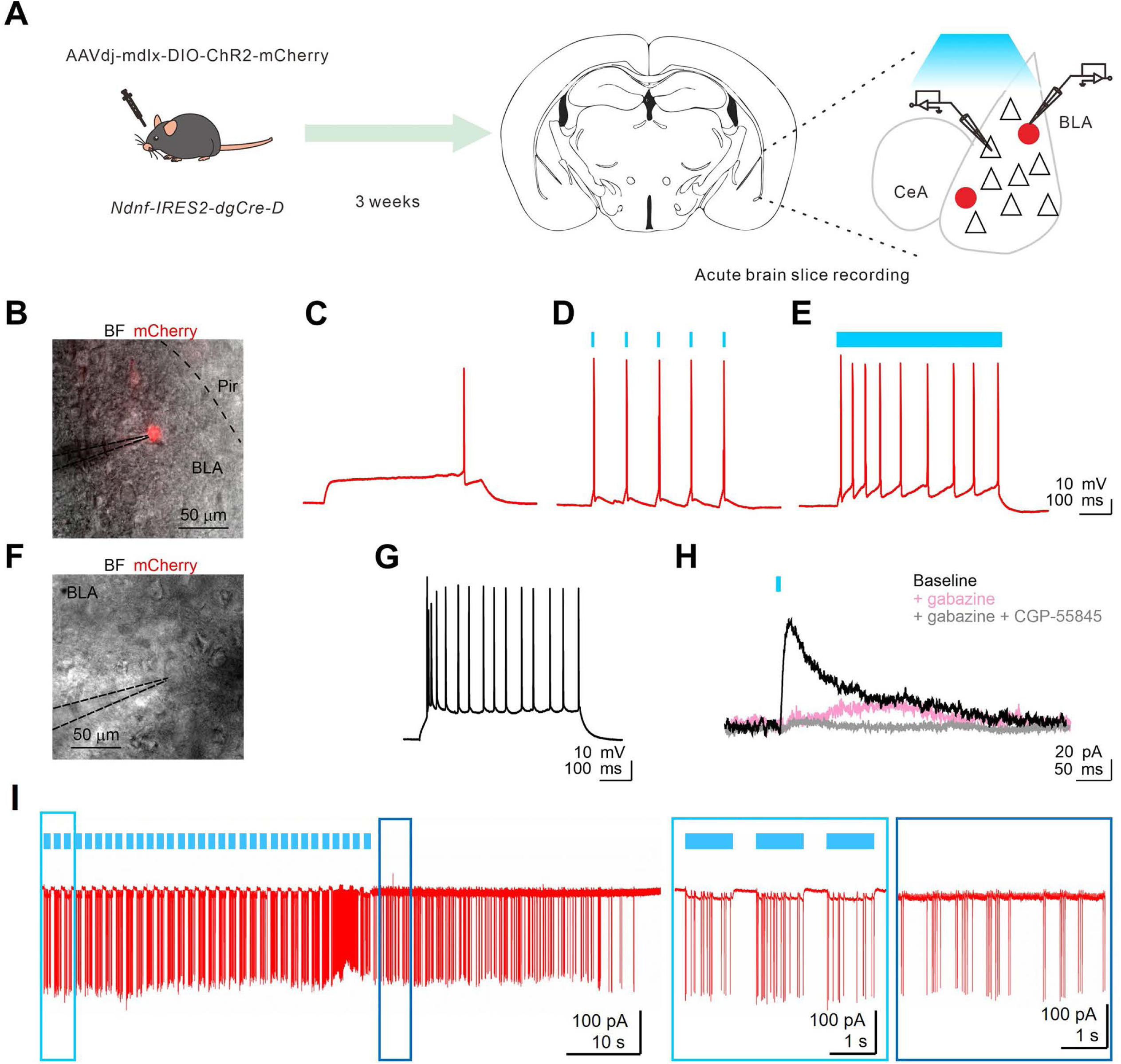
Activation of BLA NDNF+ GABAergic cells elicits both gabazine-sensitive and insensitive IPSCs. **(A)** The experimental paradigm. **(B)** An overlay image of bright field (BF) and red fluorescence signal of a representative recorded BLA mCherry+ neuron. **(C)** The rheobase response upon current injection (+80 pA, 1 s). **(D)** The membrane responses of the recorded neuron are shown on panel (C) upon short (10 Hz, 5 ms, 5 pulses) and **(E)** long pulse (1 s) photostimulation. **(F)** An overlay image of BF and red fluorescence signal of a representative recorded BLA mCherry(-) PN. **(G)** The firing pattern of the PN is shown on panel (F) upon a current injection step (+100 pA, 1 s). **(H)** PN responses to 10-Hz light pulse trains under voltage clamp configuration. Photostimulation (single pulses, 5 ms, 0.016 Hz) evoked outward currents at -60 mV (black), which could be partly blocked by bath application of GABAAR antagonist, gabazine (gabazine, 10 μM, pink). The residual slow outward current was eliminated by applying selective GABA_B_R antagonist, CGP-55845 (gray, n = 3 of 3 tested cells, n = 2 mice). No light-evoked inward currents were observed from recorded PNs throughout the optogenetic assessment (n = 10 cells, n = 5 mice). (I) Optogenetic induction of retroaxonal barrage firing with repetitive blue light stimulation (1 s, 0.6 Hz) when recording from ChR2 expressing NDNF+ neurons in the cell-attached configuration. Light and dark blue boxes show enlarged sections of triggered and retroaxonal barrage firing patterns. Abbreviations: CeA, central amygdala; Pir, piriform cortex.

After confirming successful optogenetic control over the BLA NDNF+ cell population, we performed whole-cell recordings on BLA PNs to test the potential synaptic connection. We found slow IPSCs were elicited in BLA PNs upon single pulse or 10 Hz pulse trains (10 of 10 cells, n = 5 mice; Fig.4F-H). In a subset of experiments, we further blocked GABA_A_R using gabazine (10 µM), which effectively abolished the fast component of the light-evoked PSCs, and the remaining slow outward component can be further abolished by subsequent application of selective GABA_B_R antagonist, CGP-55845 (10 µM, Fig.4H). No inward current was observed after the blockade of GABAAR-and GABABR-mediated IPSCs (3 of 3 cells, n = 2 mice; Fig.4H). These results demonstrate the optogenetic activation of BLA NDNF cell populations produces both GABA_A_R- and GABA_B_R-mediated IPSCs. Lastly, we tested if the barrage firing of BLA NDNF+ cells can be induced by optogenetic activation. In 4 out of 4 ChR2 expressing NDNF+ cells under cell-attached configuration, the optogenetic stimulation triggered the retroaxonal barrage firing (0.6 Hz, 1 s; Fig.4I).

## Discussion

The key obstacles to understanding amygdala function include the diversity of cell types, the complex connectivity, and its role in behavior. In the past decade, the advanced knowledge of developmental genetic profiling has made systematic screening and dissecting of amygdala cell types possible (Huang, 2014; Fisher and Kepec, 2019). Here, we report an approach for specifically targeting the neurogliaform cells in the mouse BLA by combining viral transfection and the NDNF-cre transgenic mouse line.

Our results indicate that a precise Cre-mediated genetic recombination could facilitate cell type investigation in the brain. The mdlx enhancer may prevent the leaky expression of Cre in the excitatory neurons of NDNF-cre mice reported in the hippocampus (Guo et al., 2021; Mercier et al., 2022). In this study, we did not detect BLA eGFP+ cells with PN-like morphology (Fig.1-2), nor did we observe any signature of evoked putative postsynaptic glutamatergic responses from recorded BLA PNs by optogenetic activating virally labeled BLA NDNF+ cells (Fig.4H). We also do not record any ChR2-mediated photocurrent directly from BLA PNs (n = 10 cells from 5 mice). These results suggest absent/undetectable Cre expression or Cre-dependent recombination occurred in BLA PNs. Other alternative approaches, such as restricting the active time window of Cre recombinase using tamoxifen-inducible Cre, may also be applied to enhance the target cell selectivity (Cre-ERT2, see Abs et al., 2018).

NGFCs in different brain areas express NDNF (Tasic et al., 2016; Poorthuis et al., 2018; Abs et al., 2018; Mercier et al., 2022). In this study, the vast majority of BLA NDNF+ cells showed similar passive and active electrophysiological properties to previously reported BLA NGFCs (Fig.2E, Table 1; Manko et al.,2012; Vereczki et al., 2021). However, a small fraction of the cases exhibit sharper AHP and faster repolarization (Fig.2D). Since we were unable to recover the anatomical features of these few cells, it remains unknown whether there are electrophysiological heterogenous NDNF+ cells or result from unspecific labeling.

**Table 1.** Summary of passive and active membrane properties of recorded BLA NDNF-GFP+ interneurons.

| Parameters | Mean ± SEM (n = 29 cells) |
| --- | --- |
| Rin (MΩ) | 238.21 ± 18.28 |
| AP width (ms) | 0.88 ± 0.05 |
| AP amplitude (mV) | 62.87 ± 1.49 |
| AHP amplitude (mV) | 13.77 ± 0.95 |
| AP adaptation index | 0.8 ± 0.072 |
| Membrane time constant (ms) | 13.67 ± 0.21 |
| Cm (nF) | 0.065 ± 0.004 |
| Sag ratio | 0.07 ± 0.01 |
| Firing frequency (Hz) | 6.9 ± 0.77 |

Abbreviations: AHP, afterhyperpolarization; AP, action potential; Cm, membrane capacitance; Rin, input resistance.

NDNF is highly enriched in the superficial layer of the neocortex and does not largely overlap with major interneuron neurochemical markers such as parvalbumin, Sst, and vasoactive intestinal peptide (Abs et al., 2018; Schumann et al., 2019). Nevertheless, the potential heterogeneity of NDNF+ GABAergic interneuron populations has also been described in the mouse somatosensory cortex as NDNF expression can be detected in GABAergic canopy cells, which are functionally distinct from NGFCs (Schumann et al., 2019). Furthermore, in non-cortical-like brain structures, such as lateral habenula, NDNF expression fails to selectively label NGFCs (Webster et al., 2021), while NGFCs derived from MGE and CGE show distinct molecular properties (Tricoire et al., 2011; Overstreet-Wadiche and McBain, 2015; Niquille et al., 2018). Further investigation is thus required to determine whether the combinatory strategy presented in this study covers the entire BLA NGFC population. Moreover, a small fraction of BLA NDNF+ cells do not express NPY nor Reln; therefore, it remains unclear whether NDNF+ cells in the BLA form a homogenous population or compose a range of NGFC types, as has been comprehensively delineated in the neocortex (Gomez et al., 2023).

Retroaxonal barrage firing is a conserved feature reported in neocortical and hippocampal areas across species. This phenomenon has predominantly been observed in NGFCs (Suzuki et al., 2014; Wang, 2015; Chittajallu et al., 2020; Rózsa et al., 2023). We found that 81% (n = 21) of BLA NGFCs exhibited persistent firing properties, indicating a high susceptibility to integrate prolonged activity in the local microcircuit. Furthermore, we validated the feasibility of eliciting persistent firing by optogenetic activation, proposing a potential application to investigate the role of BLA NGFCs’ persistent firing activity in behaviors. Persistent activity of NGFCs in the BLA could be a compensatory internal mechanism that suppresses local activity and prevents overexcitation (Suzuki et al., 2014). However, it is well-reported that NGFC synapses pose strong short-term use-dependent depression; hence, the functional relevance and effectiveness of synaptic inhibition are arguable (Tamas et al., 2003; Karayannis et al., 2010). In the BLA, ∼80% of NDNF+ GABAergic interneurons co-express NPY (Fig.1C,E). Therefore, the potential release of vasoactive substances or other molecules such as insulin (Molnar et al., 2014) during persistent firing may serve a regulatory role in fine-tuning energy and oxygen supplies in the micro-environment by increased network activity (Taube and Bassett, 2003; Echagarruga et al., 2020). However, the exact mechanisms of barrage firing, the physiological entrainment of BLA NDNF+ interneuron’s persistent firing, and its behavioral relevance under possible pathophysiological conditions deserve further investigation.

When examining the synaptic output of BLA NDNF+ GABAergic interneurons using optogenetics, we confirmed the characteristic slow inhibition from postsynaptic PNs, a feature of NGFCs described earlier (Manko et al., 2012; Tamas et al., 2003; Olah et al., 2009; Rovira-Esteban et al., 2019). Indeed, optogenetic activation is sufficient to recruit postsynaptic GABA_B_R in addition to the GABA_A_R on the target cells, contrary to any other known BLA interneuron types (Fig.4H; Woodruff and Sah, 2007; Vereczki et al., 2021).

Tonic inhibition plays a significant role in regulating the overall excitability and information processing of the brain circuitry (Marowsky et al., 2012). Such inhibition may serve as the offset control by suppressing spontaneous activity of PNs and enhancing the signal-to-noise ratio of incoming stimuli (Hausser and Clark, 1997; Krook-Magnuson and Huntsman, 2005; Semyanov et al., 2004). This fine-tuning helps to shape the population activity and facilitates the precision of temporal association of distinct salient inputs within the amygdala. A decrease in tonic inhibition can lead to hyperexcitability and altered network dynamics, which may contribute to pathological conditions like anxiety disorders or epilepsy (Tasan et al., 2011; Babaev et al., 2018; Fritsch et al., 2009). In the central amygdala, extrasynaptic inhibition is tightly linked to anxiety levels (Lange et al., 2014; Botta et al.,2015). However, the dynamics of BLA tonic inhibition in different behaviors and their causal relationships remain poorly understood. Since NGFCs are considered one of the major contributors to slow-phasic and tonic GABA release in several brain regions, including BLA (Capogna, 2014; Hajos, 2021), selective manipulation of BLA NGFC activity may help to elucidate the role of BLA tonic inhibition in distinct behaviors by future optogenetic and chemogenetic interrogations.

## Author Contributions

MC, WHH, AO, and MCS contributed to the conception and design of the study. WHH, AO, and MCS performed stereotaxic injections and acute slice electrophysiology experiments. WHH, AO, and MDS performed immunohistochemistry experiments and image acquisition. AO and WHH performed the data- and statistical analysis. WHH and AO wrote the first draft of the manuscript. MC and MDS received funds that supported this study. All authors contributed to the manuscript revision and read and approved the submitted version.

## Acknowledgments

We thank the Viral Vector Facility (VVF) of the Neuroscience Center Zurich (ZNZ) for producing the viral vectors used in this study. We also acknowledge the AU Bioimaging Core Facility and the Department of Molecular Biology and Genetics Bioimaging facility for using the equipment and the daily support from animal facilities at the Department of Biomedicine, Aarhus University. We thank Dr. Kai-Yi Wang and all the members of Marco Capogna’s lab for their comments and feedback on the manuscript.

## Data Availability Statement

The original contributions presented in the study are included in the article/supplementary material, further inquiries can be directed to the corresponding author.

